# The effect of increased CpG and UpA dinucleotides in the West Nile virus genome on virus transmission by *Culex* mosquitoes and pathogenesis in a vertebrate host

**DOI:** 10.1101/2025.01.22.634234

**Authors:** Joyce W. M. van Bree, Imke Visser, Eleanor M. Marshall, Wessel W. Willemsen, Carmen van de Waterbeemd, Marleen H. C. Abma-Henkens, Gorben P. Pijlman, Monique M. van Oers, Barry Rockx, Jelke J. Fros

## Abstract

Vertebrate animals and many small DNA and single-stranded RNA viruses that infect vertebrates have evolved to suppress genomic CpG dinucleotides. All organisms and most viruses additionally suppress UpA dinucleotides in protein coding RNA. Synonymously recoding viral genomes to introduce CpG or UpA dinucleotides has emerged as an approach for viral attenuation and vaccine development. However, studies that investigate the effects of this recoding strategy on viral replication and pathogenesis *in vivo* are still limited. Flaviviruses including West Nile virus (WNV) are transmitted between vertebrate hosts by invertebrate vectors. In humans, WNV infection can cause flu-like symptoms and neuroinvasive disease. We investigated how alterations in WNV dinucleotide frequencies impact virus replication, transmission by vector mosquitoes, as well as pathogenesis and neuroinvasiveness in vertebrates. In *Culex pipiens* vector mosquitoes and *Culex* cell lines only WNV with elevated UpA frequencies displayed attenuated replication. In vertebrate cell lines and primary human neuro-astrocyte co-cultures both UpA and CpG enrichment reduced viral replication. In mice, the CpG-high WNV mutant demonstrated partial attenuation with delayed weight loss compared to wild-type WNV, though infection still resulted in 100% mortality. In contrast, 75% of animals survived inoculation with the UpA-high WNV mutant and were protected against wild-type WNV challenge. Notably, all animals that succumbed to infection had similar levels of virus in the brain, irrespective of the WNV mutant. Our results underscore the complex interplay between viral genome composition and host immune responses, highlighting potential safety concerns for dinucleotide manipulation as a strategy for live-attenuated vaccine development in flaviviruses.

**Importance:** Flaviviruses such as West Nile virus (WNV) pose significant public health concerns due to their potential to cause severe neurological disease. Synonymously recoding flavivirus genomes to introduce CpG or UpA dinucleotides has emerged as an approach for viral attenuation and vaccine development. However, the *in vivo* effects of manipulating these frequencies across the complete transmission cycle remained unexplored. Our study provides comprehensive insights of how CpG and UpA recoding affects WNV replication in both the mosquito vector and vertebrate hosts. We demonstrate that elevated UpA content attenuates virus replication throughout the transmission cycle, while CpG enrichment only impacts replication in the vertebrate host. Although UpA-high WNV shows significant attenuation and provides protection against wild-type infection, animals that succumb exhibit similar brain viral loads as wild-type infections. These findings have critical implications for live-attenuated vaccine development based on dinucleotide manipulation, specifically highlighting the importance of carefully evaluating the risk of neuroinvasion.

## Introduction

The co-evolutionary arms race between viruses and their hosts is a dynamic process, characterized by continuous bilateral genetic adaption, including both synonymous and non-synonymous mutations. Evolutionary processes can lead to widespread biases in genome composition, such as suppression of CpG dinucleotides (cytosine followed by guanine) in vertebrates, and suppression of TpA (UpA in RNA) dinucleotides (thymine/uracil respectively, followed by adenosine) across all organisms (1, 2). The under-representation of CpG can be attributed to extensive DNA methylation, where the cytosine of a CpG dinucleotide is methylated to 5-methylcytosine (3). This can result in a methylation-induced mutation, where 5-methylcytosine deaminates to thymine (4–7). Unlike the situation in vertebrates, DNA methylation activity in invertebrate species is generally low (8, 9). Consequently, flies and mosquitoes from the order Diptera, which exhibit extremely low DNA methylation activity, demonstrate no CpG bias in dinucleotide frequency (6). The universal UpA under-representation in vertebrates and invertebrates is hypothesized to be caused by specific recognition and targeting of this dinucleotide by RNA-degrading enzymes in the cytoplasm (10).

Most viruses mimic the reduced UpA dinucleotide frequencies of their host’s mRNA, while many vertebrate infecting viruses with small DNA- and single-stranded RNA genomes additionally suppress CpG dinucleotides (11–15). This suppression of CpG dinucleotides in viral RNA most likely serves as a strategic immune evasion mechanism; by reducing CpG dinucleotide frequencies, the virus avoids recognition by the vertebrate host’s zinc-finger antiviral protein (ZAP). ZAP binds to CpG-rich single-stranded RNA sequences in the cytoplasm of vertebrate cells and can recruit additional host factors that degrade viral RNA, inhibit translation, and activate innate immune pathways, thereby suppressing virus replication (reviewed in (16)).

By introducing synonymous mutations in viral genomes, virus mutants can be created that code for wild-type viral proteins, but contain many additional CpG or UpA dinucleotides, which may expose the viral RNA to ZAP-induced antiviral responses or putative RNA degradation pathways, respectively. This approach has been explored to attenuate various RNA viruses, including human immunodeficiency virus 1 (HIV-1) (17), echovirus 7 (E7) (18, 19), influenza A virus (IAV) (20, 21) and Zika virus (ZIKV) (22, 23). Moreover, recoded IAV and ZIKV demonstrated live-attenuated vaccine potential, as mice were fully protected against a subsequent challenge with a wild-type pathogenic virus (20, 22, 23).

An interesting aspect associated with vector-borne viruses, such as flaviviruses, is their ability to replicate in evolutionarily distant invertebrate vectors (mosquitoes, ticks) and vertebrate hosts (mammals, birds), where the evolutionary pressure on CpG dinucleotides can differ (6, 22). A previous study identified opposing evolutionary pressures encountered by ZIKV when alternating between vector and host cells as the genomic introduction of CpG dinucleotides attenuated ZIKV pathogenesis in vertebrates, while it enhanced replication in *Aedes aegypti* vector mosquitoes (22). Flaviviruses are highly diverse, with distinct phylogenetic groups exhibiting variation in their primary host organisms, vector species, and mechanisms of pathogenesis. To further explore the principle that alterations in CpG and UpA dinucleotide composition impact viral replication and pathogenesis in different host species and vectors, herein we study a flavivirus that is phylogenetically distinct from ZIKV; West Nile virus (WNV). Unlike ZIKV, which primarily infects primates and is transmitted by *Aedes* spp. mosquitoes, WNV is spread by *Culex* spp. mosquitoes in a mosquito-bird transmission cycle. WNV can also infect a range of other organisms, including horses and humans, who are generally considered dead-end hosts (24). While ZIKV is primarily associated with fetal diseases, mainly targeting intermediate progenitor cells that differentiate into neurons and causing microcephaly, WNV actively replicates in the brain, leading to postnatal encephalitis and neuroinvasive disease in humans (25, 26).

In this study, we aim to assess how changes in dinucleotide frequency impact the replication, transmission and pathogenesis of WNV. We constructed synonymous WNV mutants with varying levels of CpG and UpA dinucleotides to investigate the relationship between dinucleotide composition and replication efficiency in both vertebrate hosts and the mosquito vector. We first tested the replicative phenotype of the WNV mutants in *Culex* cells and evaluated their transmission potential by *Culex pipiens* vector mosquitoes. Next we assessed their replicative fitness in vertebrate cell culture and studied replication, neuroinvasion and pathogenesis in a mouse model. Additionally, given the neuroinvasive capacity of WNV, we infected human iPCS-derived neurons to evaluate potential alterations in replication kinetics. The results of this study advance our understanding of how genomic dinucleotide frequencies influence flavivirus replication, transmission, and pathogenesis, and offers insight into the safety of manipulating CpG/UpA frequencies in WNV as a potential strategy for live-attenuated vaccine development.

## Materials and methods

### Construction of flavivirus phylogenetic tree

The following full-length coding sequences from flavivirus type species were selected from the International Committee on Taxonomy of Viruses (ICTV; https://ictv.global/): WNV: M12294, Murray Valley encephalitis virus (MVEV): AF161266, Usutu virus (USUV): AY453411, Japanese encephalitis virus (JEV): M18370, St. Louis encephalitis virus (SLEV): DQ525916, Dengue virus (DENV) 1: U88536, DENV-2: DENV-3: M93130, U87411, DENV-4: AF326573, ZIKV: AY632535, Kamiti river virus (KRV): AY149904, cell fusing agent virus (CFAV): M91671, Palm Creek virus (PCV): KC505248, Culex flavivirus (CXFV): GQ165808, Tick-borne encephalitis virus (TBEV): U27495. Multiple sequence alignment was performed using MUSCLE, and a phylogenetic tree was constructed using the Maximum Likelihood method with the General Time Reversible (GTR) model. Evolutionary history was inferred from a bootstrap consensus tree generated from 500 replicates, with branches supported by less than 50% collapsed. The percentage of replicate trees in which taxa clustered together is shown alongside the branches. Heuristic search trees were obtained using Neighbor-Joining and BioNJ algorithms, with the best topology selected based on log likelihood. Evolutionary rate differences were modelled using a discrete Gamma distribution (+G, parameter = 1.0737), with 8.98% of sites treated as invariable (+I). Analyses were conducted using MEGA X. The composition of full-length coding sequences were determined using the “composition scan” function within the Simmonds Sequence Editor (SSE) program (version 1.4) (27).

### Cell culture

Vertebrate cell lines, including African green monkey kidney Vero E6 (ATCC CRL-1586), Vero (ATCC CCL-81), human A549 cells (ATCC CCL-185) and A549 ZAP knock out (K/O) cells (28), were maintained in complete Dulbecco’s modified Eagle medium (DMEM; Gibco) supplemented with 10% (v/v) heat-inactivated fetal bovine serum (FBS; Gibco), 100 U/ml penicillin and 100 µg/ml streptomycin (pen/strep; Sigma-Aldrich). Cells were cultured at 37°C in a humidified atmosphere containing 5% CO_2_. Invertebrate cell lines, *Culex pipiens* (*C.pip*) CPE/LULS50 (*C.pip*; CVCL_C1YI) and *Culex tarsalis* (CxT) (29) cells, were cultured at 28°C in Leibovitz L-15 medium (Gibco) supplemented with 20% (v/v) FBS or Schneider’s medium (Gibco) supplemented with 10% (v/v) FBS, 2% (v/v) tryptose phosphate broth (Gibco) and 1% (v/v) non-essential amino acids (Gibco), respectively. Cell lines were passaged bi-weekly at a confluency of ∼90-100%.

### Human iPSC acquisition and culture

Human induced pluripotent stem cells (iPSCs) [WTC-11 provided by Bruce R. Conklin, the Gladstone Institutes and UCSF Institute, San Francisco, CA, USA, (UBMTA 2023-0123)] were maintained in complete iPSC medium (Stemflex medium (ThermoFisher Scientific)), 100 IU/ml penicillin (Lonza), 100 µg/ml streptomycin (Lonza) and 1x Revitacell (ThermoFisher Scientific) at 37°C and 5% CO_2_. Accutase (Life Technologies) was used for dissociation during passaging and cells were seeded on Matrigel (Corning, 10μl/ml) coated 6-well plates. For coating, Matrigel was diluted in Knock-Out Dulbecco’s Modified Eagle Medium (KO DMEM; ThermoFisher Scientific) and plates were incubated for a minimum of 1 h at 37°C. iPSCs were differentiated into spinal cord motor neurons and Ngn2 cortical neuron-astrocyte co-cultures as previously described in detail (30).

### Construction of infectious complementary DNA-clones and rescue of recombinant West Nile virus mutants

A pCC1-BAC-based infectious clone containing WNV isolate 578/10 from Hungary (lineage 2, GenBank accession no. KC496015.1) (31) was used to construct WNV mutants. Two specific regions within the WNV genome, comprising 1642 and 1645 nucleotides (nt), were selected for mutation. The scrambled control (SCR) mutant was designed with the “scramble sequence” function and the CDLR algorithm using the SSE program (27). The CpG- and UpA-elevated regions were designed using the “mutate sequence” function, adjusting CpG or UpA dinucleotide frequencies, while keeping the other dinucleotide (CpG or UpA) stable (Supplementary text 1). Regions were synthesized (GeneArt, ThermoFisher Scientific) and cloned into pCC1-BAC-WNV_578/10_ using restriction enzymes: *BstB*I and *Sph*I (region 1), and *Sph*I and *Kfl*I (region 2) (New England Biolabs). Plasmid DNA was purified using the NucleoBond Xtra Midi kit (MACHEREY-NAGEL) and subsequently transfected into Vero E6 cells using Lipofectamine 2000 (Invitrogen). Three days post-transfection, 200 µL cell culture was used to infect a ∼70% confluent monolayer of Vero E6 cells in a T-75 flask, generating passage 1 (P1) and similarly a P2 was generated. RNA from P2 was extracted using TRIzol LS Reagent (ThermoFisher Scientific) and reverse transcribed into cDNA using the SuperScript III Platinum Two-Step qRT-PCR Kit (ThermoFisher Scientific). The sequence integrity of the mutant viruses was confirmed by sequencing (Macrogen).

### Virus growth kinetics

To determine the growth kinetics of WNV, cultured cells were infected with 10 (mosquito cells) or 1 (vertebrate cells) viral RNA copies per cell. After 2 hours of incubation, the inoculum was removed, cells were washed twice with phosphate-buffered saline (PBS, Gibco), and fresh medium was added to the cells. Supernatant samples were collected daily. Infectious virus titers were subsequently determined by EPDA.

### Virus quantification

The infectious virus titer was quantified by end-point dilution assays (EPDA) on Vero cells and expressed as median tissue culture infectious dose (TCID_50_)/ml. Serial 10-fold dilutions of viral stocks and samples were prepared in complete DMEM cell culture medium, mixed in a 1:1 ratio with a Vero cell suspension of 2.5 x 10^5^ cells/ml and plated in 6-fold into microtiter plates (Nunc, Sigma-Aldrich). Viral titers were determined by the presence of cytopathic effects (CPE) 3-4 days post-infection, using the Reed-Muench method.

Viral RNA copy numbers were quantified using quantitative PCR (qPCR). Viral RNA was purified from supernatant samples using TRIzol LS. An RNA standard for WNV was generated from a 910 bp PCR amplicon corresponding to a fragment within the non-structural protein 5 (NS5) region with the T7 promoter sequence at the 5’ end (FW: TAATACGACTCACTATAGGGGAAGTGAAACCAACCGGCTCAG; RV: AGGTGTTCAGGGCGTAAGTC). RNA was *in vitro* transcribed using T7 polymerase (NEB), purified from gel (Cytiva GFX PCR and DNA Gel Band Purification Kit). Serial 10-fold dilutions of the RNA standard were prepared and used as reference for RNA quantification via SYBR Green (Fisherbrand) Real-Time quantitative PCR (RT-qPCR). The qPCR was conducted using primers targeting a 157 bp fragment within the NS5 region (FW: AAGAACGCCCGGGAAGCC; RV: TGCTGCCTTTAGCTTTGCCG) on a CFX96 Real-Time PCR instrument (Bio-Rad).

### Mosquito rearing

*Culex pipiens* mosquitoes (colony originated from Brummen, The Netherlands) were kept in Bugdorm cages and maintained at 23°C, with a 16:8 light:dark cycle and a relative humidity of 60% as described previously (32, 33).

### Infectious blood meal

Adult *Cx. pipiens* mosquitoes (5-7 days old) were starved for sucrose two days and water one day prior to the infectious blood meal. Mosquitoes were then fed with chicken whole blood (KempKip, Uden, The Netherlands) containing ∼1-5 x 10^7^ TCID_50_/ml of infectious wild-type WNV. Ratios of RNA copies between wild-type WNV and all mutants were used to calculate the required volumes of the respective virus stock for each mutant to be added to the blood. Blood feeding was conducted in a dark room for 1 h using a Hemotek PS5 feeder (Discovery Workshops, Lancashire, United Kingdom). After feeding, mosquitoes were sedated with CO_2_, and only fully engorged female mosquitoes were selected. To check for equal virus uptake, three fully engorged females of each group were immediately stored at −80°C in SafeSeal microtubes (Starstedt) containing 0.5 mm zirconium oxide beads (Next Advance). The remaining mosquitoes were incubated at 28°C for 14 days, during which they were provided with a 6% glucose solution.

### Mosquito salivation assay

Mosquitoes were sedated using CO_2_ and legs and wings were removed. The proboscis of sedated mosquitos was inserted into a 200 µL tip containing 5 µl of salivation medium (50% FBS, 25% (w/v) sucrose in dH_2_O). Mosquitoes salivated for 1 h, after which saliva was collected in Eppendorf tubes containing 55 µl complete DMEM and individual bodies in Eppendorf tubes containing 0.5 mm zirconium beads. Saliva and bodies were stored at −80°C.

### Infectivity and transmission efficiency assay

Frozen mosquito bodies were homogenized using a Bullet Blender Storm (Next Advance, New York, USA) in 100 μl of complete DMEM supplemented with 1:100 Amphotericin B (Gibco). Following homogenization, samples were centrifuged for 90 seconds at 14,000 rpm in an Eppendorf centrifuge. From the resulting supernatant and from saliva samples, 30 μl was taken and incubated on a monolayer of Vero cells in a 96-well plate. After an incubation period of 2–4 hours, the medium was replaced with 100 μl of fresh, fully supplemented medium. Three days post-infection, CPE was scored as an indication for infectivity and transmission efficiency. Viral titers from positive body samples were determined by EPDAs.

### Ethics statement mouse experiments

All animal procedures were performed in compliance with the Dutch legislation for the protection of animals used for scientific purposes (2014, implementing EU Directive 2010/63) and other relevant regulations. The licensed establishment where this research was conducted (Erasmus MC) has an approved OLAW Assurance # A5051-01. Animal experiments were conducted under a project license from the Dutch competent authority and the study protocol #174312 was approved by the institutional Animal Welfare Body.

### Intraperitoneal WNV mutant inoculation of wild-type C57BL/6 mice

Specific pathogen-free wild-type (WT) C57BL/6 age-matched female mice were obtained from Charles River Laboratories. Mice were divided into groups of 4 animals per HEPA-filtered negatively-pressured isolator filtertop-cage and allowed to acclimatize for 7 days prior to inoculation. Food and water was provided *ad libitum* throughout the duration of the experiment. All mice were 6-8 weeks old on the day of WNV inoculation. All invasive animal procedures were performed under inhalation anaesthesia using isoflurane (800-1000 ml/min O_2_, 4-5% for induction, 2-3% for maintenance). Per condition, 8 mice were inoculated intraperitoneally with a single dose (10^5^ TCID_50_) of either WNV-WT, WNV-SCR, WNV-CpG1+2, WNV-UpA1+2, or sterile PBS (control group) in a volume of 100 µl using a 25G 0.5×16mm Microlance needle (BD300600). An inoculation dose of 10^5^ TCID_50_ per animal has previously been shown to be lethal in C57BL/6 mice (34) and was therefore specifically chosen herein to test the replication, safety, and neuroinvasive capacity of the mutants. For a duration of 14 days, mice were weighed and monitored daily for general health status and behaviour and any clinical signs of disease and abnormal behaviour were documented. A 30 µl blood sample was collected from the tail vein every other day post-inoculation (up until day 10 post-inoculation) of 4/8 animals per mutant (i.e. one of both inoculated groups) and spun in serum tubes (Greiner Bio-one 450533, MiniCollect Tube 0.5 ml/0.8 ml CAT Serum Sep Clot Activator) at 4000 rpm for 10 minutes to separate and aspirate the serum. The serum was subsequently transferred to lysis buffer (Roche, MagNA Pure 96 External Lysis Buffer 06374913001) and stored at −80°C until further processing. Mice were euthanized by cervical dislocation under inhalation anaesthesia when the humane end-point was reached. Part of the spleen, liver, kidney, lung, and brain (divided into a frontal, central, and basal section) were collected in 2 ml tubes (Greiner) containing one ceramic sphere (MP Biomedicals, 11-654-0424). Tissues were stored at −80°C until further processing, then thawed once and homogenized using the Tissue Lyser II (Qiagen, 85300), and clarified by centrifugation (Allegra X-15R Centrifuge, Beckman Coulter) at 4000 rpm for 10 minutes. Subsequently, the tissue supernatants were titrated onto Vero cells as described above. Of the surviving mice, 30 µl blood was collected on day 21 post-inoculation in order to measure antibody titers and thereby confirm seroconversion of the animals.

### Intradermal WNV challenge of pre-inoculated wild-type C57BL/6 mice

Passage 3 virus stocks of the WNV-NL20 (originally isolated from a *Phylloscopus collybit*, GenBank accession number OP762595.1, EVAg 010V-04311, grown in Vero cells at 37°C in 5% CO_2_) was used for challenge. On day 28 post-inoculation, surviving mice were challenged with 10^5^ TCID50 wild-type (WT) WNV-NL20 via intradermal inoculation in the footpad of the right hind leg as described previously (35). Mice were weighed and monitored for general health status and behaviour on a daily basis and any clinical signs of disease and abnormal behaviour were documented for a duration of 10 days. A 30 µL blood sample was collected from the tail vein every other day post-challenge (up until day 10 post-challenge) and processed as described above. On day 10 post-challenge, or when the humane end-point was reached, mice were euthanized and tissues were harvested as described above.

### Serum RNA isolation and real-time qPCR

Total RNA was extracted from the serum samples by transferring the serum samples to a 96-well plate and incubation with Agencourt AMPure XP magnetic beads (Beckman Coulter A63882) for at least 15 minutes. Samples were then placed onto a DynaMag-96 magnet (Invitrogen, 12027) for 3 minutes, after which the supernatant was aspirated and discarded. The beads were washed three times with 70% ethanol and air-dried for 3 minutes while kept on the magnetic block. RNA was eluted from the beads by resuspension in sterile ddH2O. Real-time TaqMan PCR was performed using a WNV primer/probe mix (Probe sequence TGCTGCTGCCTGCGGCTCAACCC, Forward: CCACCGGAAGTTGAGTAGACG, Reverse: TTTGGTCACCCAGTCCTCCT) diluted in TaqMan Fast Virus-1 Step Master Mix (Applied Biosystems 4444432) and ddH2O. The Applied biosystems 7500 Real-Time PCR system (ThermoFisher Scientific) was used with the following program settings: 5 minutes 50°C, 20 seconds 95°C, 45 cycles of 3 seconds 95°C and 30 seconds 60°C. A serial dilution of WNV-578/10 or WNV-NL20 viral stock for the generation of a standard curve was included for comparison and conversion to TCID_50_ equivalents per ml post-mutant and challenge inoculation respectively.

### Virus titrations of mouse tissues

Viral titers (expressed as the median tissue culture infectious dose; TCID_50_/ml) were determined by an EPDA, titrating supernatant on Vero cells seeded onto 96-well plates at a density of 2.3×10^4^ cells per well ∼24 hours prior to titration. The inoculated plates were incubated at 37°C in 5% CO_2_, after which cytopathic effects were assessed 6 days post-infection (days post-infection) to determine the virus titers using the Spearman & Kärber method (36).

### Statistical analysis

All statistical analyses were conducted using GraphPad Prism software (version 10.2.3). Differences in CpG and UpA O/E ratios between *Culex* and *Aedes* transmitted vertebrate-infecting flaviviruses were tested using an unpaired Student’s T-test. To compare replication kinetics between the mutants and WT WNV, total viral load was calculated by calculating the total area under curve (AUC). Differences in AUC were analyzed using the non-parametric Friedmann test, followed by Dunn’s multiple comparison. A non-parametric Kruskal-Wallis test with Dunn’s multiple comparison was used to check for significant differences in blood titers and body titers between WT WNV or the other WNV variants containing blood or infected mosquitoes. For the mouse experiment, statistical analyses were done using a 2-way ANOVA multiple comparison to assess significant differences in weight loss and viremia, and a log-rank Mantel-Cox test to assess differences in mortality.

## Results

### Dinucleotide suppression patterns in flaviviruses correlates with host and vector specificity

To further explore the relationship between dinucleotide frequencies, host range, vector species and putative phylogenetic groups within the *Orthoflavivirus* genus, we constructed a concise phylogenetic tree (Fig. 1). The phylogeny shows that flaviviruses cluster into distinct groups that are defined by vector species, host tropism and the pathogenesis they induce. Adjacent to the phylogenetic tree, we placed color-codes representing the observed-over-expected (O/E) ratios of CpG and UpA dinucleotides within the full-length genomes of each virus. Vertebrate-infecting flaviviruses exhibit a marked suppression of both CpG (O/E = 0.38 – 0.62) and UpA (O/E = 0.43 – 0.60) dinucleotide frequencies, while insect-specific flaviviruses only show suppression of UpA dinucleotides (O/E = 0.49 – 0.66).

**Figure 1.**
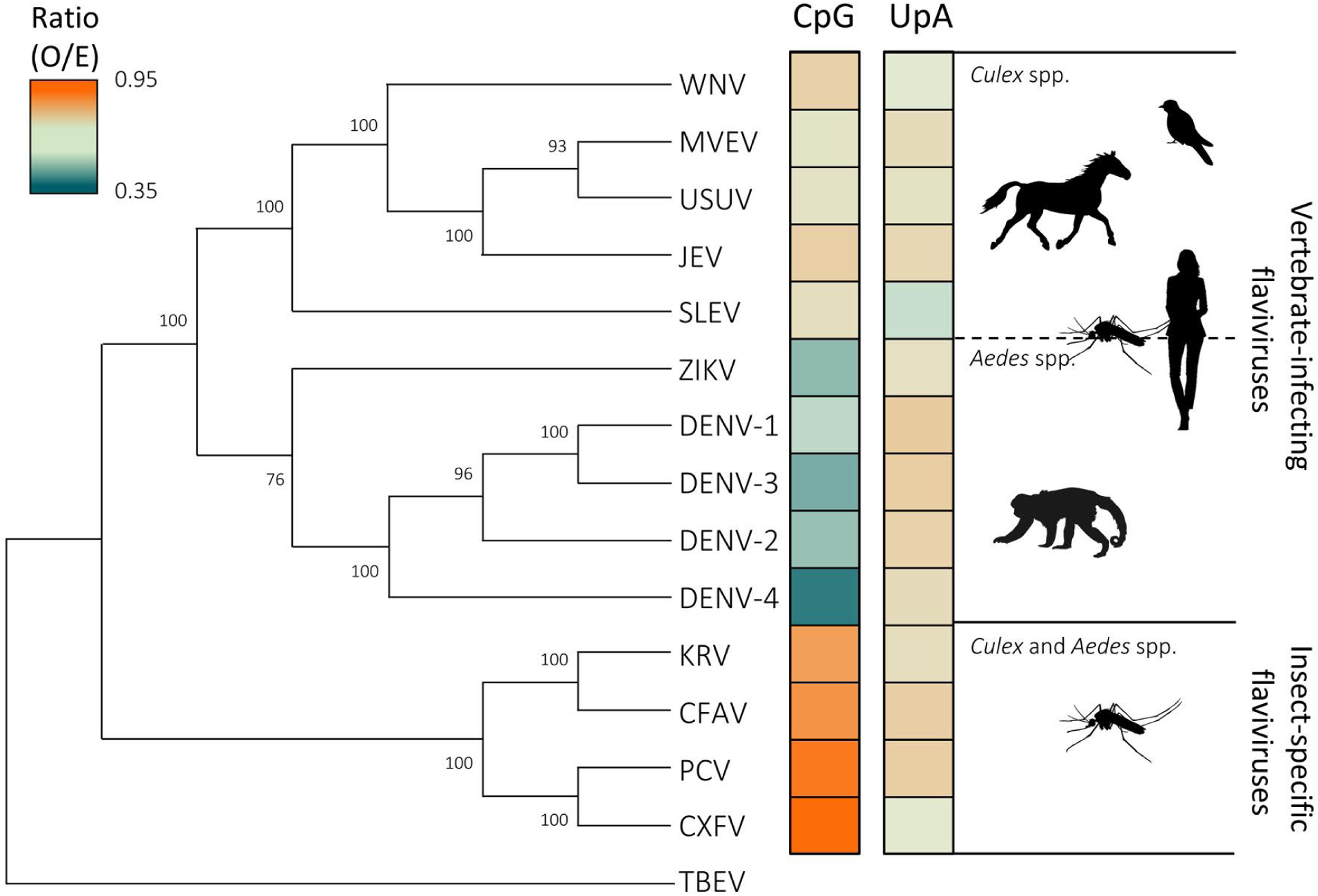
Phylogenetic relationships and dinucleotide composition of the genus *Orthoflavivirus*. A phylogenetic tree of selected viruses from the genus *Orthoflavivirus* (West Nile virus (WNV): M12294, Murray Valley encephalitis virus (MVEV): AF161266, Usutu virus (USUV): AY453411, Japanese encephalitis virus (JEV): M18370, St. Louis encephalitis virus (SLEV): DQ525916, Dengue virus (DENV) 1: U88536, DENV-3: M93130, DENV-2: U87411, DENV-4: AF326573, Zika virus (ZIKV): AY632535, Kamiti river virus (KRV): AY149904, cell fusing agent virus (CFAV): M91671, Palm Creek virus (PCV): KC505248, Culex flavivirus (CXFV): GQ165808, Tick-borne encephalitis virus (TBEV): U27495). TBEV was used for rooting the tree. The color-codes adjacent to the tree represent the observed-to-expected (O/E) ratios of CpG and UpA dinucleotides within the genomes of each virus. Evolutionary history was inferred from a bootstrap consensus tree generated from 500 replicates, with branches supported by less than 50% collapsed. Bootstrap values are displayed alongside branches. Heuristic search trees were obtained using Neighbor-Joining and BioNJ algorithms, with the best topology selected based on log likelihood. Evolutionary rate differences were modelled using a discrete Gamma distribution (+G, parameter = 1.0737), with 8.98% of sites treated as invariable (+I). CpG suppression was significantly stronger in *Aedes*-transmitted flaviviruses (O/E = 0.42) compared to *Culex*-transmitted viruses (O/E = 0.55; *p* < 0.001, unpaired Student’s t-test). UpA O/E ratios showed no significant difference between both transmission groups (*p* > 0.05).

Interestingly, in this phylogenetic display, we also observed differences in CpG suppression between vertebrate-infecting flaviviruses transmitted by *Aedes* and *Culex* mosquitoes, with CpG-suppression being significantly (*p* < 0.001) more pronounced in *Aedes*-transmitted flaviviruses (O/E = 0.42) compared to those transmitted by *Culex* mosquitoes (O/E = 0.55). The differences between ZIKV and WNV in host species, vector preference, and CpG O/E ratios (0.43 vs 0.58) led us to investigate how dinucleotide alterations affect WNV replication and transmission.

### Construction of West Nile virus mutants with increased CpG and UpA dinucleotides

To investigate how CpG and UpA dinucleotides affect replication and pathogenesis of a Culex-transmitted, neuroinvasive flavivirus, various synonymous WNV mutants were designed. First, genomic regions were selected that could accommodate synonymous mutations without disrupting essential functional elements. The flavivirus genome contains key RNA structures in the 5’ and 3’ untranslated regions (UTRs) involved in genome cyclization (reviewed in Ng et al., 2017), and the conserved heptanucleotide motif and pseudoknot in the NS2A region, required for ribosomal frameshifting and NS1’ production (38, 39). Remaining genomic regions display relatively high and uniform nucleotide variability at synonymous sites (40), that are potentially suitable sites to introduce synonymous mutations. Two regions in the WNV genome were selected for introducing additional CpG or UpA dinucleotides: region 1, coding for the polyprotein from NS3 to NS4A (1642 nt), and the adjacent region 2, coding for NS4A to NS5 (1645 nt) (Fig. 2A).

**Figure 2.**
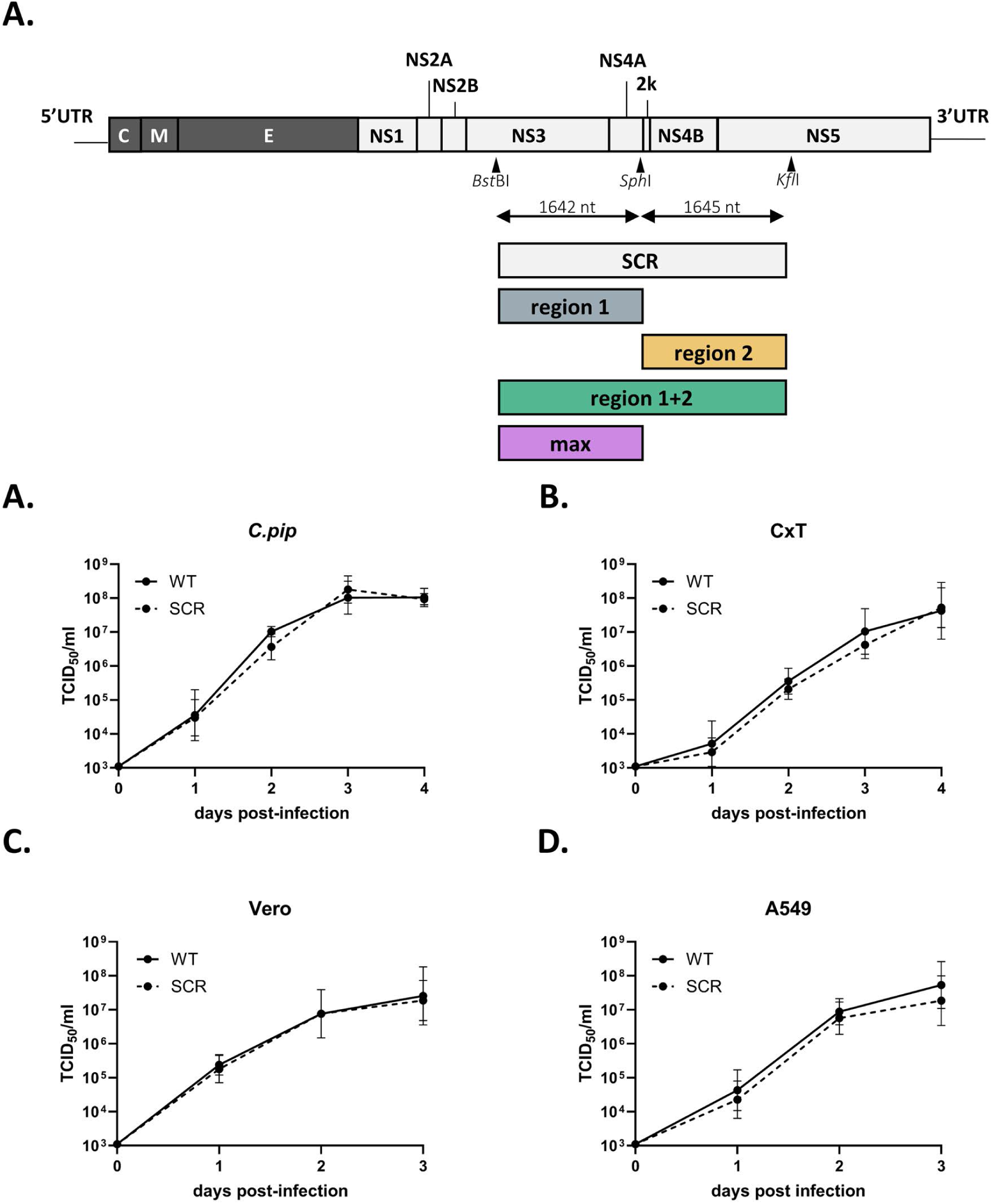
West Nile virus mutant design and scrambled control replication kinetics. (A) Selected genomic regions for synonymous mutations introducing CpG and UpA dinucleotides in the West Nile Virus (WNV) genome. Replication of wild-type (WT) and scrambled (SCR) WNV in (A) *C.pip* and (B) CxT cells at 0, 1, 2, 3 and 4, and (C) Vero and (D) A549 cells at 0, 1, 2, and 3 days post-infection. Viral titers in cell supernatants were quantified over time. Data points represent the mean of three independent experiments, and error bars show the standard error of the mean. Differences between viral titers were not significant (Friedman test with Dunn’s multiple comparison *p* > 0.05).

Second, a scrambled control (SCR) mutant was constructed. This SCR mutant incorporates the maximum number of synonymous changes within the target regions while preserving the wild-type mono- and dinucleotide frequencies. The SCR mutant virus exhibited replication kinetics similar to wild-type WNV across both mosquito and vertebrate cell lines, as measured by infectious progeny virus in the cell culture supernatant (Fig. 2B-E). These results confirmed that the selected regions lack essential RNA structures or genomic sequences, making them suitable for testing the effects of CpG and UpA dinucleotide modifications on WNV replication.

We then generated mutants with elevated CpG or UpA frequencies. In region 1, 49 additional CpG dinucleotides or 57 additional UpA dinucleotides were introduced to achieve unbiased observed/expected (O/E) ratios of approximately 1.0. Similarly, 47 CpG or 51 UpA dinucleotides were added in region 2 to reach the same target ratio (Table 1). In these mutants, the final mononucleotide frequencies were identical to the wild-type sequence. Additionally, in CpG mutants, the UpA frequencies were kept at wild-type levels and *vice versa* to isolate the effects of each modification.

**Table 1.** Synonymous CpG and UpA dinucleotide modifications in West Nile virus mutants. Wild-type (WT) and scrambled control (SCR) mutants maintain baseline dinucleotide frequencies, while CpG and UpA mutants exhibit targeted increases in CpG or UpA dinucleotides. The CpG and UpA observed/expected (O/E) ratios indicate the relative frequency of these dinucleotides within each mutant genome.

|  | Length (bp) | G+C (%) | CpG<sup>#</sup> | CpG (O/E) | UpA<sup>#</sup> | UpA (O/E) |
| --- | --- | --- | --- | --- | --- | --- |
| WT full genome | 10302 | 51.0 | 379 | 0.57 | 280 | 0.46 |
| WT1+2 | 3287 | 51.5 | 117 (+0) | 0.54 | 80 (+0) | 0.42 |
| SCR | 3287 | 51.5 | 117 (+0) | 0.54 | 80 (+0) | 0.42 |
| CpG1 | 1642 | 50.4 | 103 (+49) | 0.99 | 40 (+0) | 0.41 |
| CpG2 | 1645 | 52.5 | 110 (+47) | 0.98 | 40 (+0) | 0.43 |
| CpG1+2 | 3287 | 51.5 | 213 (+96) | 0.99 | 80 (+0) | 0.41 |
| CpGmax | 1642 | 60.2 | 150 (+96) | 1.25 | 40 (+0) | 0.64 |
| UpA1 | 1642 | 50.4 | 54 (+0) | 0.52 | 97 (+57) | 0.99 |
| UpA2 | 1645 | 52.5 | 63 (+0) | 0.56 | 91 (+51) | 0.99 |
| UpA1+2 | 3287 | 51.5 | 117 (+0) | 0.54 | 188 (+108) | 0.99 |
| UpAmax | 1642 | 50.4 | 54 (+0) | 0.52 | 139 (+99) | 1.42 |

<sup>#</sup>Total number of introduced CpG or UpA dinucleotides displayed as (+N)

To assess the cumulative impact of dinucleotide enrichment, combined mutants were created incorporating modifications from both regions. It was also investigated whether the spatial distribution and concentration of these dinucleotides affect viral replication and attenuation by constructing CpGmax and UpAmax mutants. These variants contained concentrated numbers of dinucleotides in region 1 equivalent to the cumulative number introduced in regions 1+2 combined, with 96 CpG or 99 UpA dinucleotides, respectively (Table 1). For the CpGmax variant this was only achievable when mononucleotide usage was allowed to change from WT levels, as reflected in the slightly elevated G+C content (Table 1).

### Replication of West Nile virus mutants with increased CpG and UpA dinucleotides in mosquito cells

To evaluate the impact of CpG- and UpA-enrichment on WNV replication in *Culex* mosquito vector cells, *Culex pipiens* (*C.pip*) and *Culex tarsalis* (CxT) cell cultures were infected with the WNV variants and infectious progeny virus was measured. For the CpG-high mutants, overall replication kinetics were not significantly different compared to WT WNV in both cell lines (Fig. 3A+B). In contrast, the UpA-high mutants exhibited consistently reduced viral titers compared to WT WNV in both *C.pip* and CxT cells (Fig. 3C+D). Notably, a statistically significant reduction in viral titers was observed for the UpA2 in both cell lines (*p* < 0.05). These findings suggest that the CpG content has no effect on WNV replication, while increasing the UpA content results in lower WNV titers in *Culex* mosquito cells.

**Figure 3.**
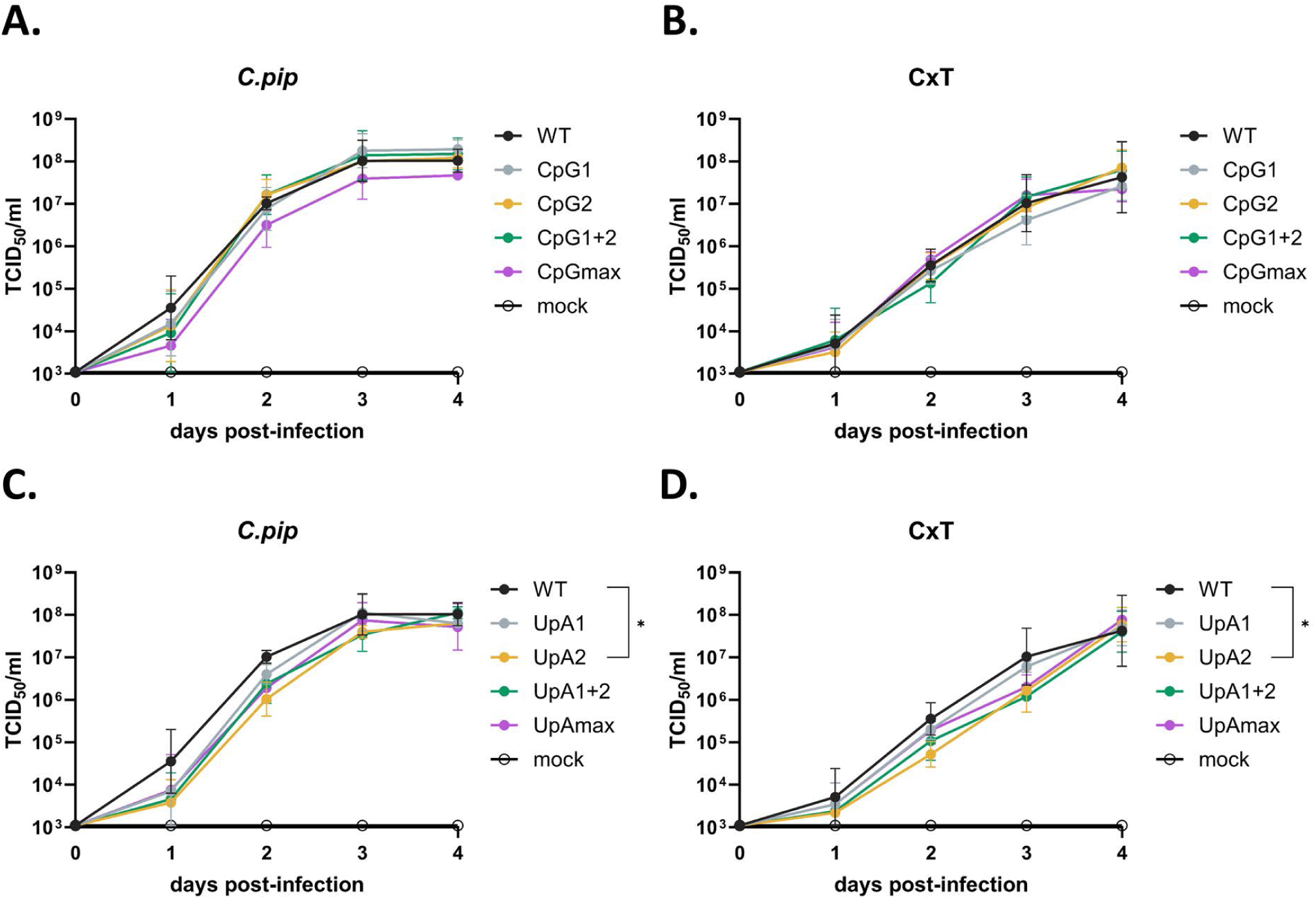
Replication kinetics of West Nile Virus CpG- and UpA-high mutants in mosquito cells. Cells were infected at a multiplicity of infection (MOI) of 10 RNA copies/cell. (A) Replication of CpG mutants and (B) UpA mutants in *C. pipiens* (*C.pip*) cells, measured at 0, 1, 2, and 3 days post-infection. (C) Replication of CpG mutants and (D) UpA mutants in *Cx. tarsalis* (CxT) cells under identical conditions. Viral titers in cell supernatants were quantified over time. Data points represent the mean of three independent experiments, and error bars show the standard error of the mean. Statistical analysis on the area under curve for each virus variant was performed by using the non-parametric Friedman test with Dunn’s multiple comparisons. Asterisks indicate significant differences ((*) *p* < 0.05).

### Replication of West Nile virus mutants with increased CpG and UpA dinucleotides in *Culex pipiens* mosquitoes

To investigate the replicative phenotypes of WNV mutants in live vector mosquitoes, *Culex pipiens* were orally infected with either WT, SCR, Region 1 (R1), Region 1+2 (R1+2) and the max (Rmax) CpG and UpA mutant viruses (Fig 2A). All variants were delivered at a fixed dose of 10^10^ RNA copies/ml, corresponding to ∼2 x 10^7^ TCID_50_/ml for WT WNV. Blood meal and three fully engorged female mosquitoes were collected immediately after feeding to determine exposure titers per group. Blood containing CpGmax WNV showed slightly lower titers, and viral titers in mosquito bodies fed with CpGmax WNV were reduced compared to WT WNV (Supplementary Fig. 1).

Following infection, mosquitoes were incubated and monitored for 14 days. Survival was high across all WNV variants (above 96%), except for WNV-CpGmax, which had a slightly lower survival rate of 88% (Fig. 4A). Among surviving mosquitoes, saliva was harvested before bodies were homogenized and analyzed for the presence WNV. Infectivity rates, measured as the proportion of mosquitoes with detectable virus in body homogenates, were highest in the WT WNV group (65%) and lower in the mutants. The lowest infectivity rate was observed in the CpG1 group (31%), followed by CpGmax (40%) and UpAmax (41%) (Fig. 4B). Notably, despite reduced infectivity, CpGmax-infected mosquitoes exhibited high median viral titers (median = 5.6) in positive samples, nearly matching those of WT-infected mosquitoes (median = 6.2) (Fig. 4C). Median body titers in UpA1 and UpAmax-infected mosquitoes were significantly lower than those in WT-infected mosquitoes (*p* < 0.05, Fig. 4C). Transmission rates, indicated by virus in saliva, followed trends similar to infectivity rates. WT-infected mosquitoes had the highest transmission rate (59%), while CpG1+2-infected mosquitoes showed the lowest (36%), followed by CpG1 and UpAmax (37%) (Fig. 4D).

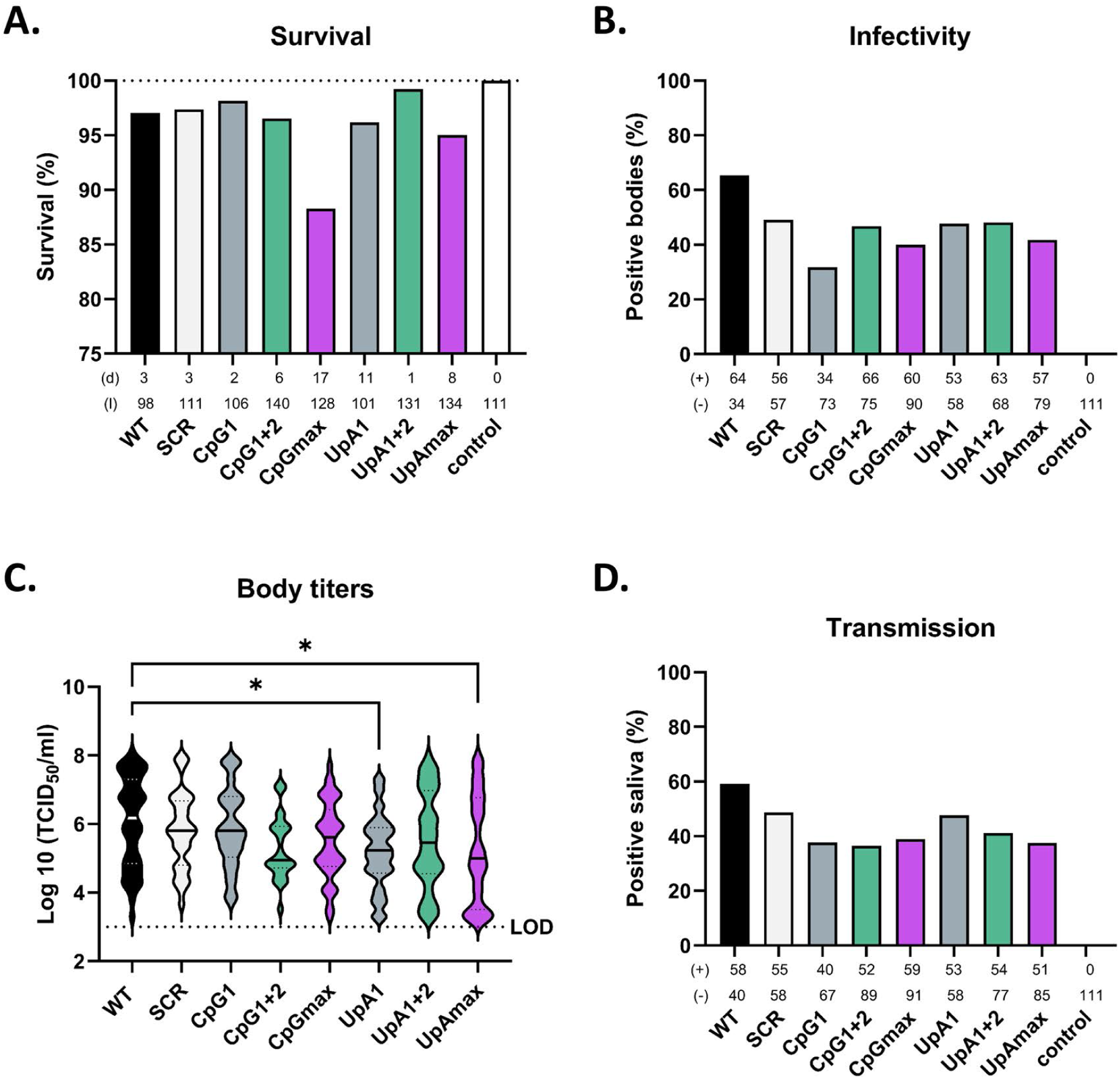

In conclusion, both infectivity and transmission were reduced across all mutants, including the SCR, suggesting that synonymous changes in selected regions may affect viral fitness *in vivo*. Despite reduced infectivity rates, CpG1 and CpGmax mutants maintained relatively high median viral titers comparable to WT WNV, while infection with UpA-high mutants resulted in lower viral titers. These findings suggest that virus replication within the mosquito, but not initial infection, is affected by elevating CpG and UpA frequencies in WNV.

### West Nile virus mutants with increased CpG and UpA dinucleotides are attenuated in vertebrate cells

To investigate the replication kinetics of WNV mutants with increased CpG and UpA dinucleotides in vertebrates, common cell lines from primate and human origin (Vero and A549, respectively) were infected with 1 RNA/cell and daily cell culture supernatant samples were collected to assess the presence of infectious virus.

In both Vero and A549 cells, the CpG-high mutants replicated slower than WNV WT, with the CpGmax mutant displaying a significant difference from WT (*p* < 0.001, Fig. 5A,C). In A549 ZAP K/O cells, the replication kinetics of all CpG mutants were similar to WT WNV, indicating that the attenuation of CpG mutants is restored in the absence of ZAP (Fig. 5E).

**Figure 5.**
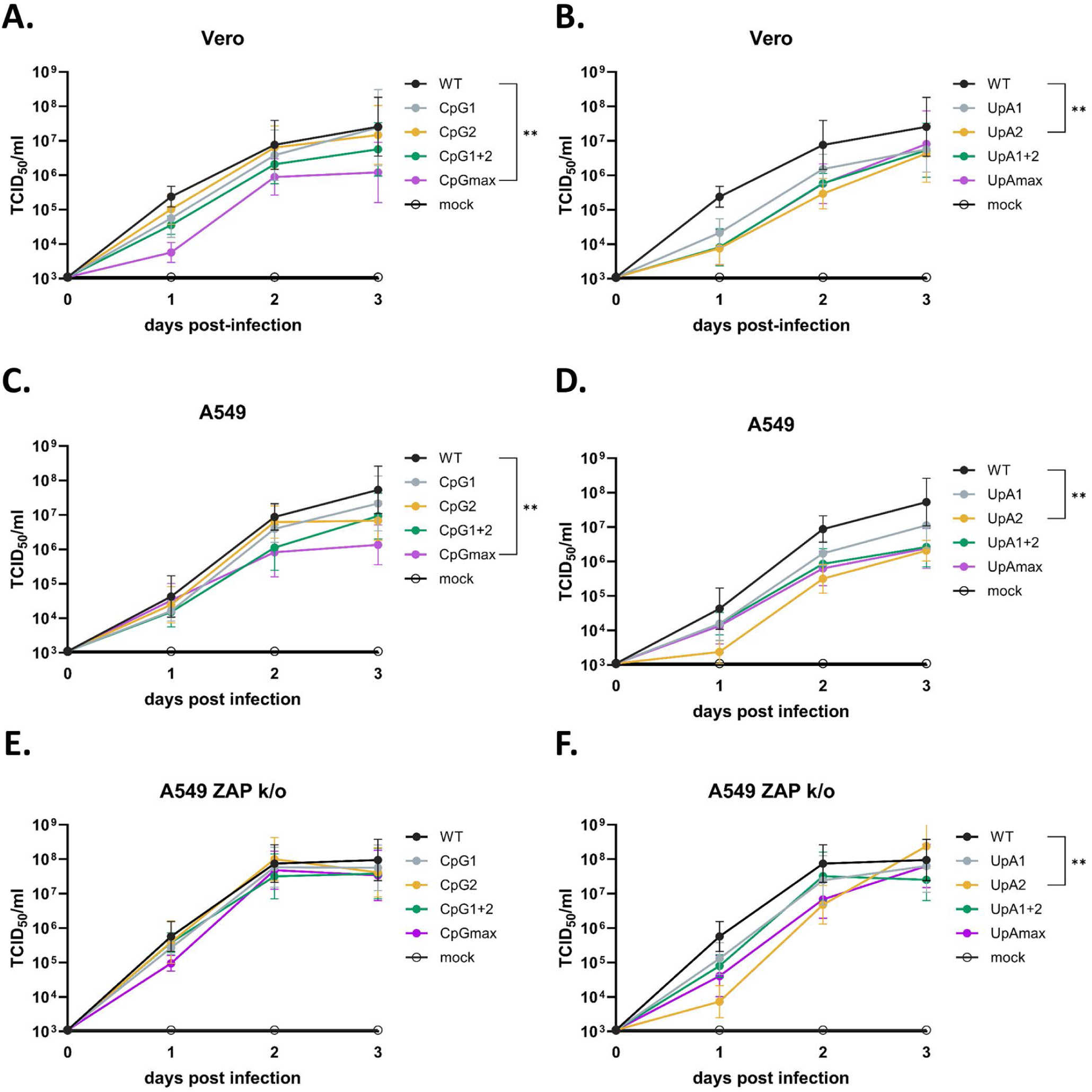
Replication kinetics of West Nile Virus CpG and UpA high mutants in vertebrate cell lines. (A+B) Vero, (C+D) A549, and (D+E) A549 ZAP knockout (ZAP k/o) cells were infected at a multiplicity of infection (MOI) of 1 RNA/cell with wild-type (WT) West Nile virus (WNV) and either (A+C+E) CpG-high mutants or (B+D+F) UpA-high mutants. Viral replication was assessed by measuring viral titers (TCID50/ml) in cell supernatants at 0, 1, 2, and 3 days post-infection. Data points represent the mean of three independent experiments, and error bars indicate the standard error of the mean. Statistical analysis on the area under curve for each virus variant was performed by using the non-parametric Friedman test with Dunn’s multiple comparisons. Asterisks indicate significant differences ((*) *p* < 0.05, (**) *p* < 0.01).

Similarly, all UpA mutants replicated to lower titers compared to WT WNV in both Vero and A549 cells, with the UpA2 mutant displaying a significant difference from WT in both cell lines (*p* < 0.001, Fig. 5B+D). Unlike the CpG mutants, the replication kinetics of the UpA mutants were not restored to WT levels in A549 ZAP K/O cells (Fig. 5F). However, titers for the different UpA-mutants were a little closer to WT WNV. These results indicate that genomic CpG and UpA dinucleotides negatively affect WNV replication in vertebrate cells.

### West Nile virus CpG and UpA mutants are attenuated in human iPCS-derived cortical-neuron astrocyte cocultures

Previous research has shown that some live-attenuated WNV strains can still replicate in the central nervous system, raising concerns about the possibility of neurovirulence (41, 42). Therefore, we evaluated the replication of WT, CpG1+2 and UpA1+2 mutant WNV strains in human iPSC-derived motor neurons and Ngn2 cortical neuron-astrocyte cocultures as a proxy for their ability to replicate within the peripheral and central nervous systems, respectively. The CpG1+2 and UpA1+2 mutants, were selected as they contain the most additional dinucleotides without altering mononucleotide freqencies (Table 1).

In the motor neurons, the replication of WT WNV was highest compared to the SCR, CpG1+2 and UpA1+2 mutants, at all time points (Fig. 6A). A significant difference was observed between WT WNV and the UpA1+2 mutant (*p* < 0.05, Fig. 6A). In neuron-astrocyte co-cultures, both CpG1+2 and UpA1+2 mutants exhibited significantly lower replication compared to WT (*p* < 0.01 and *p* < 0.05, Fig. 6B). These observations indicate that the CpG1+2 and UpA1+2 mutants are attenuated in their capacity to replicate in neurons, and that this attenuation is more apparent in cortical neuron-astrocyte co-cultures compared with motor neurons alone.

**Figure 6.**
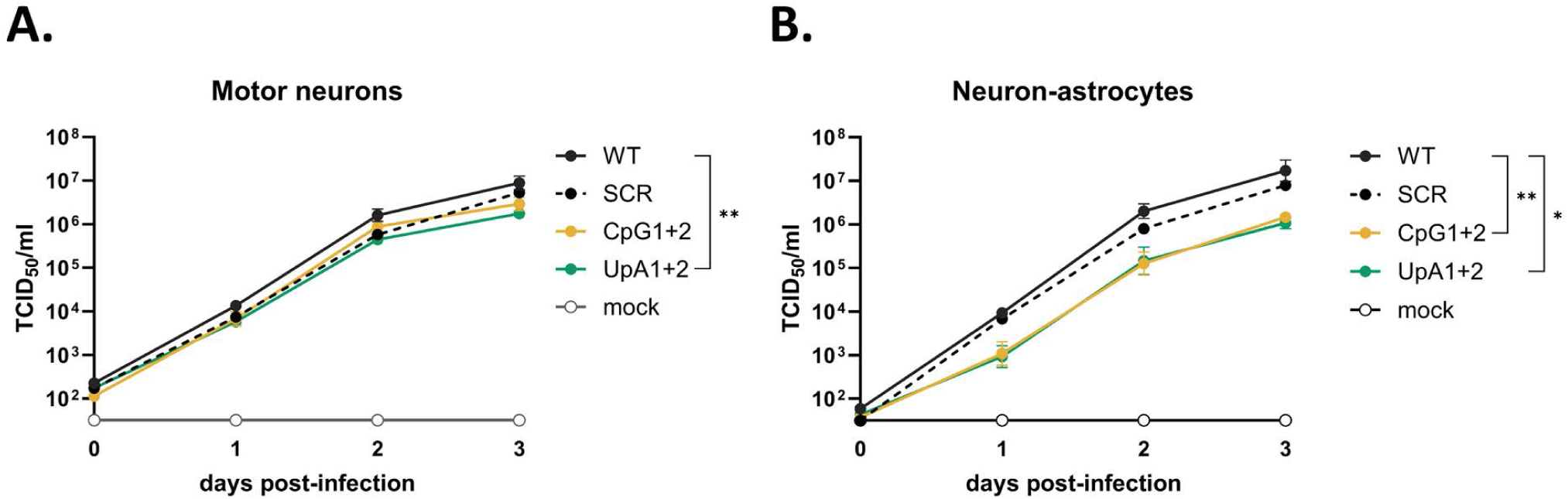
Replication kinetics of West Nile Virus CpG- and UpA-high mutants in iPCS-derived cortical-neuron astrocyte cocultures. Cells were infected at a multiplicity of infection (MOI) of 1 RNA copy/cell. (A) Replication of CpG-high and (B) UpA-high mutants in induced pluripotent stem cell-derived motor neurons, measured at 2 days post-infection. (C) Replication of CpG-high and (D) UpA-high mutants in neuron-astrocyte co-cultures at 2 days post-infection. Viral titers in cell supernatants were quantified at 0, 1, 2, and 3 days post-infection. Data points represent the mean of three independent experiments, and error bars show the standard error of the mean. Statistical analysis on the area under curve for each virus variant was performed by using the non-parametric Friedman test with Dunn’s multiple comparisons. Asterisks indicate significant differences ((*) *p* < 0.05, (**) *p* < 0.01).

### WNV-CpG and WNV-UpA induce weight loss and mortality in C57BL/6 WT mice

To assess attenuation of the WNV mutants *in vivo*, C57BL/6 WT mice were inoculated intraperitoneally with WNV-SCR, WNV-CpG1+2, or WNV-UpA1+2 and disease progression was compared to WNV-WT infection (Fig. 7A). Mock animals were injected with PBS as negative control. WT-inoculated mice started to lose weight from 5 days post-inoculation (dpi), and continued to lose weight up until 8-9 dpi, where all WT-inoculated animals reached the humane end-point (set at 75% of the starting bodyweight). Animals inoculated with CpG1+2 started losing weight at 6 dpi and despite losing significantly less weight compared to WT (Fig.7B), they all reached the humane end-point between 7-10 dpi.. In the UpA1+2-inoculated group 25% of the group started losing weight at 6 dpi, while 75% either remained relatively stable or recovered and returned to their starting weight (Fig. 7B; Supplementary Fig. 2). All animals in the WT, SCR, and CpG1+2 inoculated groups reached the humane end-point before the end of the experiment. However, the mortality of the CpG1+2 group was significantly delayed by 1 and 2 days compared to WT and SCR, respectively (Fig. 7C). In contrast, with 75% the overall probability of survival of UpA1+2 inoculated animals was significantly higher than WT-inoculated animals. These findings demonstrate that UpA1+2 WNV is more effectively attenuated than CpG1+2 compared to WT.

**Figure 7.**
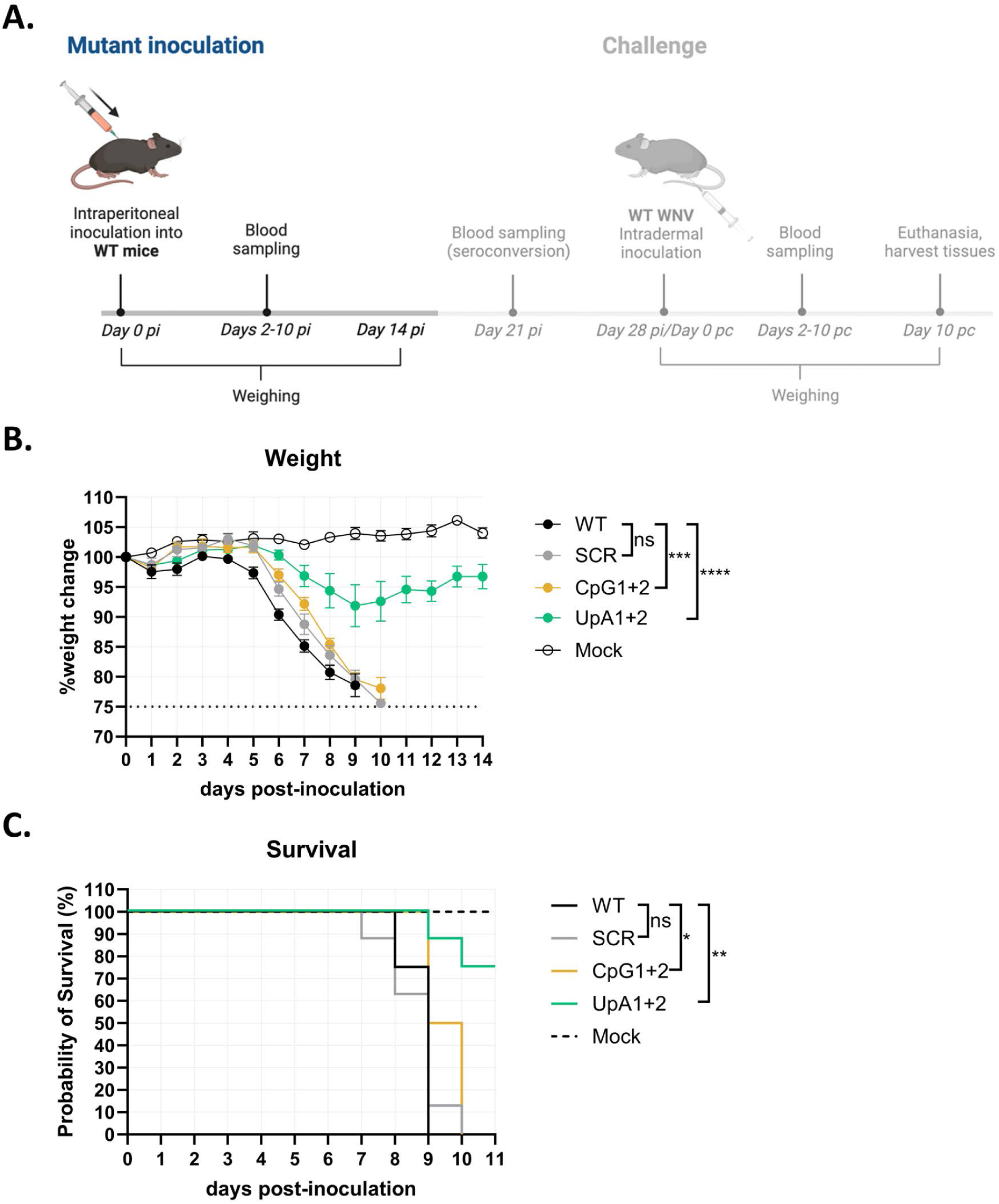
Weight loss and mortality of C57BL/6 wild-type mice post-inoculation with West Nile virus mutants. (A) Animal trial timeline for West Nile virus (WNV)-NL20 challenge in C57BL/6 wild-type (WT) mice pre-inoculated with WNV mutants. On day 0, animals were inoculated with WNV WT or mutant (10^5^ TCID_50_/mL) via the intraperitoneal route (*n* = 8 per group with the exception of WT group; *n* = 4). All animals were monitored for clinical signs of infection and weighed every day up until day 14 post-infection (pi). Blood was sampled every other day. (B) Weight change post WT, scrambled (SCR), CpG1+2, UpA1+2, and mock inoculation. Error bars represent the standard error of the mean. Statistically significant differences at day 7 pi were assessed using a 2-way ANOVA multiple comparisons test over day 0-7 pi (i.e. before the animals started to reach humane end-point). (C) Survival curves of inoculated animals. Statistical analysis was conducted using the Log-rank (Mantel-Cox) test. Asterisks indicate significant difference ((*) *p* < 0.05, (**) *p* < 0.01, (***) *p* < 0.001, or (****) *p* < 0.0001) between the corresponding mutant and WT WNV.

### WNV-CpG and WNV-UpA inoculation results in viremia and leads to detectable virus in the brain of C57BL/6 WT mice

To assess whether inoculation with the WNV mutants leads to the development of viremia, serum was collected every other day starting at 2 dpi. Viremia peaked at 2 dpi for all mutants, where both CpG1+2- and UpA1+2-inoculated animals reached significantly lower peak titers (∼10^4^ TCID_50_ /ml) compared to WT-inoculated animals (∼10^5^ TCID_50_/ml). Between 2-8 dpi, viremia gradually decreased for all mutants, where no significant difference between titers was observed between the groups (Fig. 8A). We further determined the viral load in the main visceral target organs (spleen, liver, kidney, lung) and in the brain (divided into frontal, central, and basal regions) which had been collected on the days when animals reached the humane end-point post-inoculation. No positive titers were found in the visceral organs, apart from one animal in the WT (Fig. 8B) and SCR (Fig. 8C) groups (∼10^3^ TCID_50_/g in kidney and lung). In contrast, all WT, SCR, CpG1+2 inoculated animals (Fig. 8B-D), and the two UpA1+2 (Fig. 8E) inoculated animals that had reached the humane end-point had measurable viral titers in the brain ranging from 10^3^ to 10^8^ TCID_50_/g on the day of euthanasia.

**Figure 8.**
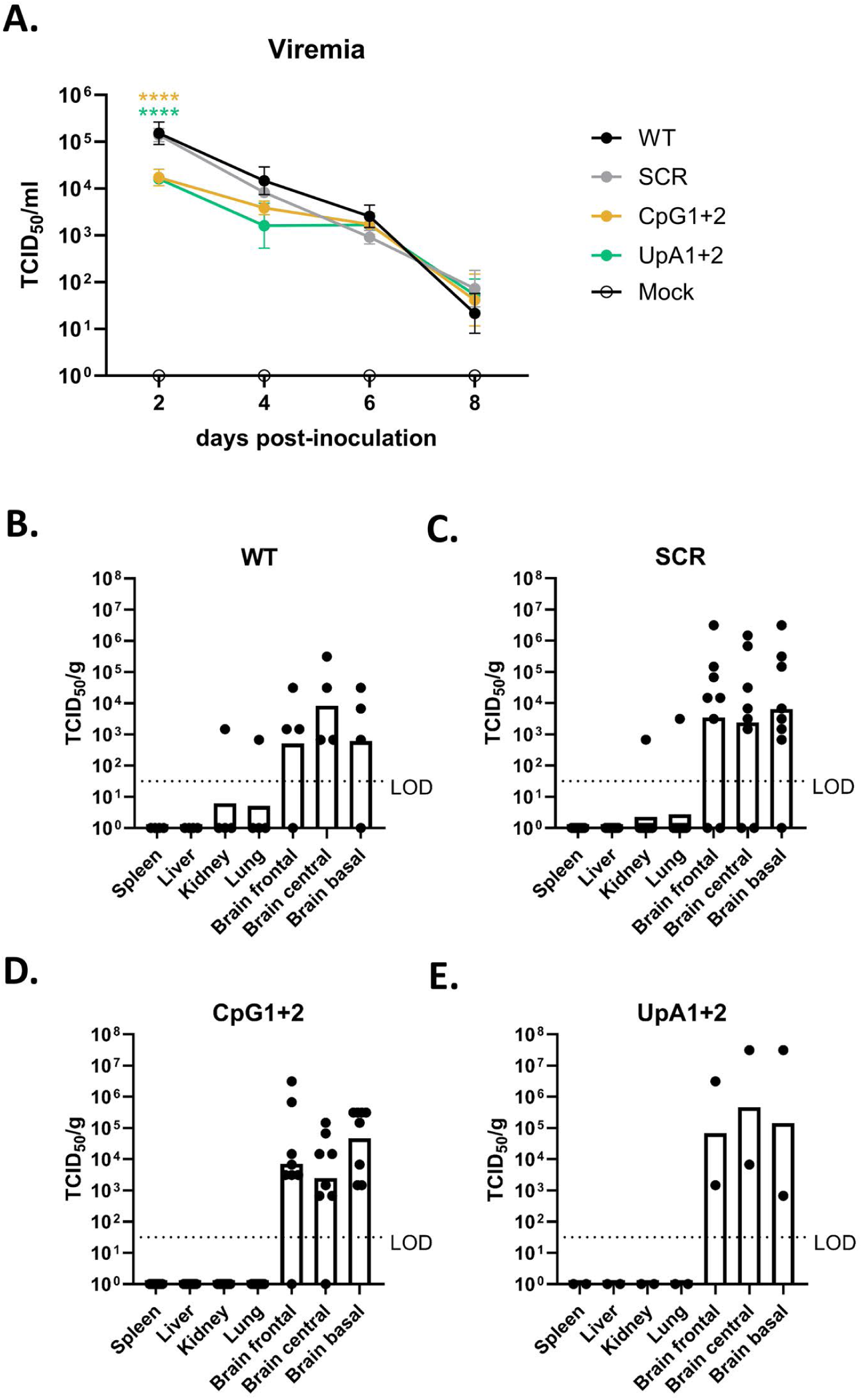
Viremia and viral loads in the blood and brains of C57BL/6 wild-type mice. (A) Viral RNA titers in the blood as determined by qPCR. Blood was sampled every 2 days post-infection up until 8 dpi (*n* = 4 per mutant), or when the humane end-point was reached. Error bars represent the standard error of the mean. Significant differences in viremia were assessed using a 2way ANOVA multiple comparison. (B-E) Infectious virus titers in different visceral organs and regions of the brain in (B) wild-type (WT)-inoculated animals (*n* = 4); (C) scramble (SCR)-inoculated animals (*n* = 8); (D) CpG1+2-inoculated animals (*n* = 8); and (E) UpA1+2-inoculated animals (*n* = 2) on the day of euthanasia, as measured by virus titration. Shown as TCID_50_ per gram of tissue. LOD = limit of detection. Asterisks and colors indicate significant difference ((*) *p* < 0.05, or (**) *p* < 0.01) between the corresponding mutant and WT WNV for that specific time point.

### WNV-UpA1+2-inoculated animals are protected against subsequent WNV-WT challenge

The surviving 75% of WNV-UpA1+2-inoculated animals all seroconverted (Supplementary Fig. 3). Therefore, to assess whether initial inoculation of the WNV-UpA1+2 could confer protection against subsequent WNV-WT infection, all remaining UpA1+2-inoculated animals (*n* = 6) were challenged with WNV-NL20 (a related WT WNV lineage 2 strain) on day 28 post-inoculation (UpA1+2-WT group). The mock-animals that had been injected with PBS previously (*n* = 8) were subsequently divided into two groups of 4 animals, where one group was challenged with WNV-NL20 (WT group) and the other group injected with PBS to again serve as a mock group (Fig. 9A).

**Figure 9.**
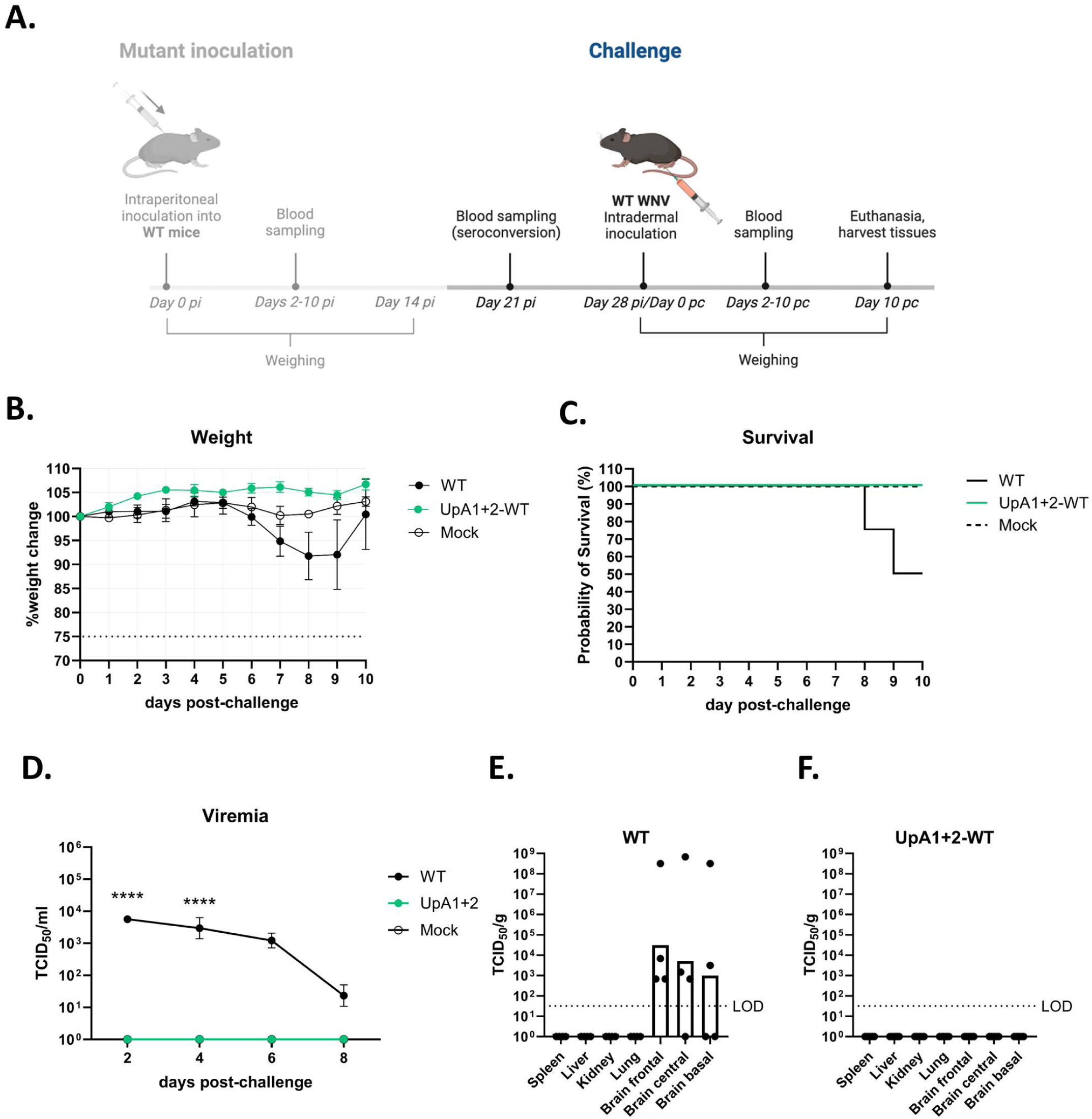
Weight loss, mortality, viremia, and tissue titers of naïve and UpA1+2 pre-inoculated animals challenged with wild-type West Nile virus. Animals were challenged intradermally in the footpad of the right hind leg with 10^5^ TCID_50_ per animal (wilde-type (WT) and mock group *n* = 4; UpA1+2v group *n* = 6). (A) Animal trial timeline for West Nile virus (WNV)-NL20 challenge in C57BL/6 wildtype mice pre-inoculated with WNV mutants (Fig. 7A). On day 21 pi, blood was collected from all animals that survived the inoculation to check for seroconversion. On day 28 pi, or day 0 post-challenge (pc), animals were challenged with the wildtype WNV-NL20 strain and monitored in the same way as pi. (B) Weight change of WT, UpA1+2, and mock animals followed over a time-course of 10 days post-challenge. Error bars represent the standard error of the mean. (C) Survival curves of challenged animals. (D) Viral RNA titers in the blood as determined by qPCR. Blood was sampled every 2 days post-challenge up until day 8, or when the humane end-point was reached. Error bars represent the standard error of the mean. Significant differences in viremia were assessed using a 2way ANOVA multiple comparison and indicated by the asterisks ((****) *p* < 0.0001). (E-F) Infectious virus titers in indicated visceral organs and regions of the brain on the day of euthanasia, as measured by virus titration, in (E) naïve animals challenged with WNV; (F) UpA1+2 pre-inoculated animals challenged with WNV. Shown as TCID_50_ per gram of tissue. LOD = limit of detection.

None of the WNV-NL20 challenged animals that had been pre-inoculated (‘vaccinated’) with UpA1+2 lost weight, in contrast to the previously naïve group that started to lose weight starting at 6 days post-WNV-NL20 challenge (Fig. 9B). Correspondingly, the probability of survival of the UpA1+2-WT inoculated mice was 100% upon challenge, compared to 50% for the WT group (Fig. 9C). In addition, UpA1+2-WT animals did not develop viremia (Fig. 9D) or tissue titers (Fig. 9F) post-challenge. All animals from the WT group developed titers in the brain (Fig. 9E). These results show that once animals survive initial inoculation with WNV-UpA1+2, they are subsequently protected against WNV disease.

## Discussion

In this study, we engineered several WNV variants with varying levels of increased CpG and UpA frequencies to better understand the evolutionary pressures that these dual-host flaviviruses encounter and whether introducing these mutations has potential for live-attenuated vaccine development to protect against neuroinvasive WNV. We evaluated the replicative fitness of WNV mutants in mosquito and vertebrate cells, assessed how the introduced changes in dinucleotide ratios influence transmission by vector mosquitoes and whether these mutations affect replication and neurovirulence in a mouse model.

In mosquitoes, the genomic frequencies of CpG dinucleotides are unbiased (6, 15). Correspondingly, the genomes of insect-specific flaviviruses exhibit no or very limited CpG suppression (43, 44). This suggests that mosquitoes do not exert an evolutionary pressure to reduce viral CpG dinucleotides. Our *in vitro* experiments showed that the replication kinetics of CpG-high mutants in *Culex* cell lines were similar to those of WT WNV (Fig. 3A+B).

When testing the replication of our mutants in *Culex* mosquitoes, we observed that CpGmax-infected mosquitoes contained relatively high viral titers and showed the highest mortality. However, these mosquitoes paradoxically exhibited a lower infectivity and transmission rate compared to the WT infected groups (Fig. 4). Our previous report showed enhanced replication and higher infectivity of CpG-high ZIKV mutants in *Aedes* cells and mosquitoes (22). Because CpG-suppression is more pronounced in *Aedes*-transmitted flaviviruses compared to those transmitted by *Culex* mosquitoes (Fig. 1), the discrepancies with our previous study may suggest that (the strength of) evolutionary pressures on CpG dinucleotides differ between *Aedes* and *Culex* mosquito vectors or viral species.

UpA dinucleotides are suppressed in the genomes of all mosquitoes and flaviviruses. Our data demonstrates that increasing UpA content attenuates WNV replication in mosquito cells, before reaching the same peak end titers as wild-type WNV. Also in live *Culex* mosquitoes, we observed an effect on infectivity, body titers and transmission between WT WNV and the UpA mutants infected groups (Fig. 4B-D). Similar results were observed with UpA-high ZIKV, showing attenuated replication in *Aedes* cells (22). Altogether, this suggests that UpA enrichment reduces the replication efficiency of flaviviruses in the mosquito vector.

In vertebrate cell lines, the number of CpGs introduced correlated to the attenuated phenotype, with consistent but less pronounced attenuation of the CpG1 and CpG2 mutants and stronger attenuation of tyhe CpG1+2 and CpGmax mutants compared to WT WNV (Fig. 5A+B). Notably, despite containing the same total number of genomic CpG dinucleotides, the CpGmax mutant exhibited greater attenuation than the CpG1+2 mutant in both Vero and A549 cells. This suggests that not only the quantity but also the spacing of CpG dinucleotides plays a critical role in WNV replication (45). In vertebrate cell lines, the UpA mutants exhibited attenuated replication compared to WT WNV. The UpA1 mutant was less attenuated compared to the UpA2, UpA1+2, and UpAmax mutants (Fig. 5D+E). Interestingly, the UpA2 mutant displayed similar levels of attenuation to the UpA1+2 and UpAmax mutants, despite differences in UpA dinucleotide frequencies, suggesting that the replicative fitness of distinct UpA mutants is differentially influenced by factors beyond dinucleotide frequency alone.

Replication of CpG-high mutants was fully restored in A549 ZAP K/O cells, confirming the central role of ZAP in CpG-mediated restriction (Fig. 5C). UpA-high mutants also showed improved replication in ZAP-knockout cells (Fig. 5F), which is in line with previous research on ZIKV and echovirus 7 (22, 28). A direct interaction between ZAP and UpA dinucleotides seems unlikely, given ZAP’s structural specificity for RNA targets containing CpG dinucleotides (46). Therefore, the partial rescue of UpA mutants in ZAP-K/O cells suggests overlapping molecular pathways in CpG and UpA recognition, though the precise mechanism remains to be elucidated.

The differential replication patterns of CpG1+2 and UpA1+2 WNV variants between motor neurons and cortical neuron-astrocyte co-cultures reveal important insights into the replication efficiency of modified viral genomes in different cell types. The similar replication kinetics between WNV variants (with only slight attenuation of UpA1+2) suggest that CpG and UpA modifications have minimal impact on viral replication in motor neurons (Fig. 6A). ZAP’s antiviral activity against CpG-high viruses requires co-factors (e.g. TRIM25) and is intertwined with innate immunity (Reviewed in (16)). It is therefore imaginable that attenuation of CpG- and UpA-high WNV may depend on innate immune activation, which WNV effectively evades in motor neurons (30). In contrast, the marked attenuation of both CpG- and UpA-high WNV variants in cortical neuron-astrocyte cultures, compared to robust WT and SCR replication (Fig. 6B), may indicate a crucial role of astrocytes in detecting modified viral genomes through their higher expression of pathogen recognition receptors and more robust innate immune responses (47, 48), or neuronal subtype specific differences in antiviral responses between cortical and motor neurons (49).

In mice, the CpG1+2 mutant displayed significantly delayed weight loss and mortality compared to WT WNV, consistent with previous findings on CpG-high IAV and ZIKV mutant infection in mice (21–23), and suggesting a modest attenuation of this mutant. Nevertheless, infection of mice with the CpG1+2 variant resulted in 0% probability of survival and equivalent viral brain titers compared to the WT WNV, which was not observed for IAV (21) and ZIKV (22) CpG-high variants. This difference in disease outcome could be due to the fact that in our study, a lethal dose of WNV was injected to specifically test the replication, safety, and potential neuroinvasiveness of the mutants.

In contrast, infection with the UpA1+2 variant resulted in clear, albeit heterogeneous, attenuation of weight loss and mortality (Fig. 7B+C). However, the two UpA1+2-infected animals that did not survive showed brain titers comparable to those observed in CpG1+2 and WT WNV-infected animals (Fig. 8B-E), indicating that once the mutant viruses overcome the initial stages of infection, they are able to reach and replicate within the brain similar to WT levels. Unfortunately, in the surviving UpA1+2-infected animals we could not determine whether the UpA1+2 mutant also successfully entered the brain or, if present, how it was subsequently cleared. To conclusively elucidate these mechanisms, a serial-sacrifice study would be necessary in order to have access to and assess (brain) samples during the acute phase of the infection. Other CpG and UpA-recoded virus species have been shown to elicit strong inflammatory responses (18, 21), leading to attenuation and full protection against subsequent WT infection in mice (21–23). The UpA1+2-infected mice that survived the initial infection (Fig. 7C), were completely protected against subsequent challenge with WT WNV (Fig. 9).

Although previous research has explored the vaccine potential of CpG-recoded viruses, our findings introduce critical nuances for such vaccine development strategies. The attenuation of recoded viruses appears to be substantially influenced by host immune competence, as exemplified by the age-dependent antiviral responses reported by Trus et al. (23). While both CpG and UpA mutants showed attenuation in neuron-astrocyte co-cultures, both mutants still reached relatively high titers (10^6^ TCID_50_/ml). Their ability to replicate in motor neuron cells and their detection in our mouse brains challenges their potential as live-attenuated vaccine candidates. This neurotropism, combined with our *in vivo* findings, highlights the need for careful evaluation of recoded WNV as a live-attenuated vaccine, particularly for individuals with compromised immune systems. However, we identified a dose- and spacing-dependent effect of CpG dinucleotides on virus replication, suggesting that the number, type, and distribution of mutations can be strategically optimized to enhance safety. Moreover, developing combination mutants with both increased CpG and UpA content may provide an additional approach to improving attenuation and safety profiles.

## Acknowledgements

This work is part of the research programme One Health PACT with project number 109986, which is (partly) financed by the Dutch Research Council (NWO) and has received funding from the European Union’s Horizon 2020 research and innovation programme under grant agreement No 952373. The work was further supported by the Dutch Ministry of Infrastructure and Water Management (31181038) and JJF by a VIDI grant from the Dutch Research Council (NWO; VI. Vidi. 213.027). Finally, we thank Felicity Chandler for performing the protein microarray; Vincent Vaes, Lars Vermaat, Ingeborg van Middelkoop, Rianne Stam, and Josianne Theuns-van Vliet for their assistance during the animal trials; and Corinne Geertsema for her technical support.

---

**Supplementary Figure 1. Blood and body titers post feeding.** (A) Viral titers in the blood meal fed to *Culex pipiens* mosquitoes, showing back-titration of the infectious blood meal containing each West Nile virus (WNV) variant. The blood meal was standardized to ∼2×10^7^ TCID50/ml for wild-type (WT) WNV to ensure consistent viral input across all groups. (B) Body viral titers of fully engorged *Culex pipiens* mosquitoes immediately after feeding. Data points represent data of three independent experiments. Error bars represent the SEM. Statistical analysis was conducted on the mean viral titers using the non-parametric Kruskal-wallis with Dunn’s multiple comparisons test. Asterisks indicate significant difference (*p* < 0.001) between the corresponding mutant and WT WNV. LOD = limit of detection.

**Supplementary Figure 2. Weight loss of WNV-UpA1+2-inoculated animals,** followed over a time-course of 14 days. Shown as weight change per animal.

**Supplementary Figure 3. Seroconversion of WNV-UpA1+2-inoculated animals**. On day 21 post-inoculation, blood was collected from all animals that had survived the mutant inoculation to check for IgG-binding titers on microarray. Signal for all UpA1+2 surviving animals was saturated (reaching the upper limit of detection).

